# Male and female gametophytes meet after winter: Testing the overwintering hypothesis for delayed fertilization in Japanese stone oak

**DOI:** 10.1101/2024.10.16.618622

**Authors:** Takenori Shagawa, Kota Ogawa, Masahiro M Kanaoka, Akiko Satake

## Abstract

**Premise:** Fertilization delays of nearly a year have been observed in many acorn-producing Fagaceae trees. Despite this long-standing recognition, the underlying mechanisms and adaptive significance of delayed fertilization remain largely unexplored. It has been hypothesized that these prolonged delays may serve as an adaptive strategy, allowing ovules to be fertilized and seeds to develop during more favorable seasons, thereby avoiding the adverse conditions of winter. However, empirical evidence to support this hypothesis is still lacking.

**Methods:** To test the overwintering hypothesis, we observed the seasonal progression of pollen tube growth and ovule development in *Lithocarpus edulis*, a species exhibiting two flowering seasons, spring and autumn. We conducted monthly observations of pollen tube growth and ovule development in both spring and autumn flower samples. We employed microtome techniques and scanning confocal microscopy to closely examine the seasonal progression of pollen tube growth and ovule development.

**Results:** We found that pollen tubes were arrested at the style-joining site, and ovules remained immature in both spring and autumn samples prior to winter. Following winter, pollen tube regrowth and ovule maturation were synchronized in the subsequent spring in both spring and autumn samples.

**Conclusions:** Our results support the overwintering hypothesis, suggesting that delayed fertilization may serve as an adaptive strategy to ensure fertilization in spring, overcoming the reproductive challenges posed by cold winter conditions. Our study introduces a new avenue for studying the temporal organization of plant reproductive phenology, specifically the alignment of flowering, pollination, fertilization, and fruiting in seasonal environments.

---

Fertilization at the appropriate timing is crucial for the reproductive strategy of plants. In angiosperms, fertilization generally takes place within 24–48 hours following pollination (Williams, 2008). However, delays in fertilization extending beyond 4 days have been reported in various taxonomic groups (Sogo and Tobe, 2006). The time lapse between pollination and fertilization can range from several days to nearly a year (Sogo and Tobe, 2006). Although the phenomenon of delayed fertilization was first reported over a century ago (Benson, 1894), the underlying mechanism and its adaptive significance remain largely unknown.

The sperm competition hypothesis proposes that the adaptive significance of delayed fertilization lies in enhancing competition among pollen tubes and enabling selective choice by the pistils for superior sperm (Sogo and Tobe, 2006; Bochenek and Eriksen, 2011). However, this hypothesis alone does not sufficiently explain the extended delay in fertilization, which can be nearly a year. As an alternative hypothesis, Satake and Kelly (2021) proposed the overwintering hypothesis. Their mathematical model, which incorporates survival and competition among female flowers, predicts that in the presence of an unsuitable season for reproduction, such as winter, a strategy evolves to delay fertilization until a more favorable season, such as spring. This model suggests that even plants flowering in spring evolve to delay fertilization for nearly a year rather than proceeding with immediate fertilization when seed maturation cannot be completed before the onset of an unfavorable season. While this hypothesis is intriguing, empirical trials to test this prediction have yet to be conducted.

In this study, we empirically tested the overwintering hypothesis by observing the process from pollination to fertilization in *Lithocarpus edulis* (Makino) Nakai. *L. edulis* exhibits two flowering seasons, spring and autumn, depending on the region (Satake et al., 2023). The family Fagaceae, one of the most diverse tree families distributed across northern temperate regions (Manos and Stanford, 2001), includes a remarkable number of species that exhibit delayed fertilization (Satake and Kelly, 2021a). In the genus *Lithocarpus*, most species mature acorns in the next year following flowering and pollination (2-year fruiting; Satake and Kelly, 2021a; Araye et al., 2022). By comparing the pollination-to-fertilization process in pistils produced in different seasons, we aim to determine whether fertilization occurs while avoiding winter. If the adaptive significance of delayed fertilization is solely explained by sperm competition, pistils produced in spring and autumn should exhibit fertilization at different timings after a certain period following flowering, irrespective of the presence of winter. Conversely, if pistils produced in both spring and autumn exhibit synchronous fertilization after winter, this would support the overwintering hypothesis. Here we tested the hypothesis by using microtome techniques and scanning confocal microscopy to observe the seasonal progression of pollen tube growth and ovule development.

## MATERIALS AND METHODS

### Plant materials and sampling methods

Female inflorescences of *L. edulis*, planted at the Ito Campus (33°35’45.3” N 130°12’42.2” E) of Kyushu University (Fukuoka, Japan), were collected monthly from July 2022 to October 2023. For each monthly sampling, one individual was chosen from eight mature trees arbitrary, and at least three inflorescences were collected from the individual. At the Ito Campus, *L. edulis* exhibits two distinct flowering periods each year, with a major flowering period in June (spring flowering) and a minor flowering period from September to October (autumn flowering; Fig. 1; Satake et al., 2023). The samples collected from spring flowering were referred to as the spring samples, while those collected from autumn flowering were designated as the autumn samples. The female inflorescences for the spring samples were collected from June of the first year to June of the subsequent year, while those for the autumn samples were collected from November of the first year to June of the subsequent year (Fig. 1C). The inflorescences were immediately fixed in FAA (70% ethanol: stock formalin: acetic acid, 90:5:5, by volume). The samples were transferred to the laboratory within 2 hr after sampling and stored at 4 °C for at least 48 hours until further investigation.

**Figure 1.**
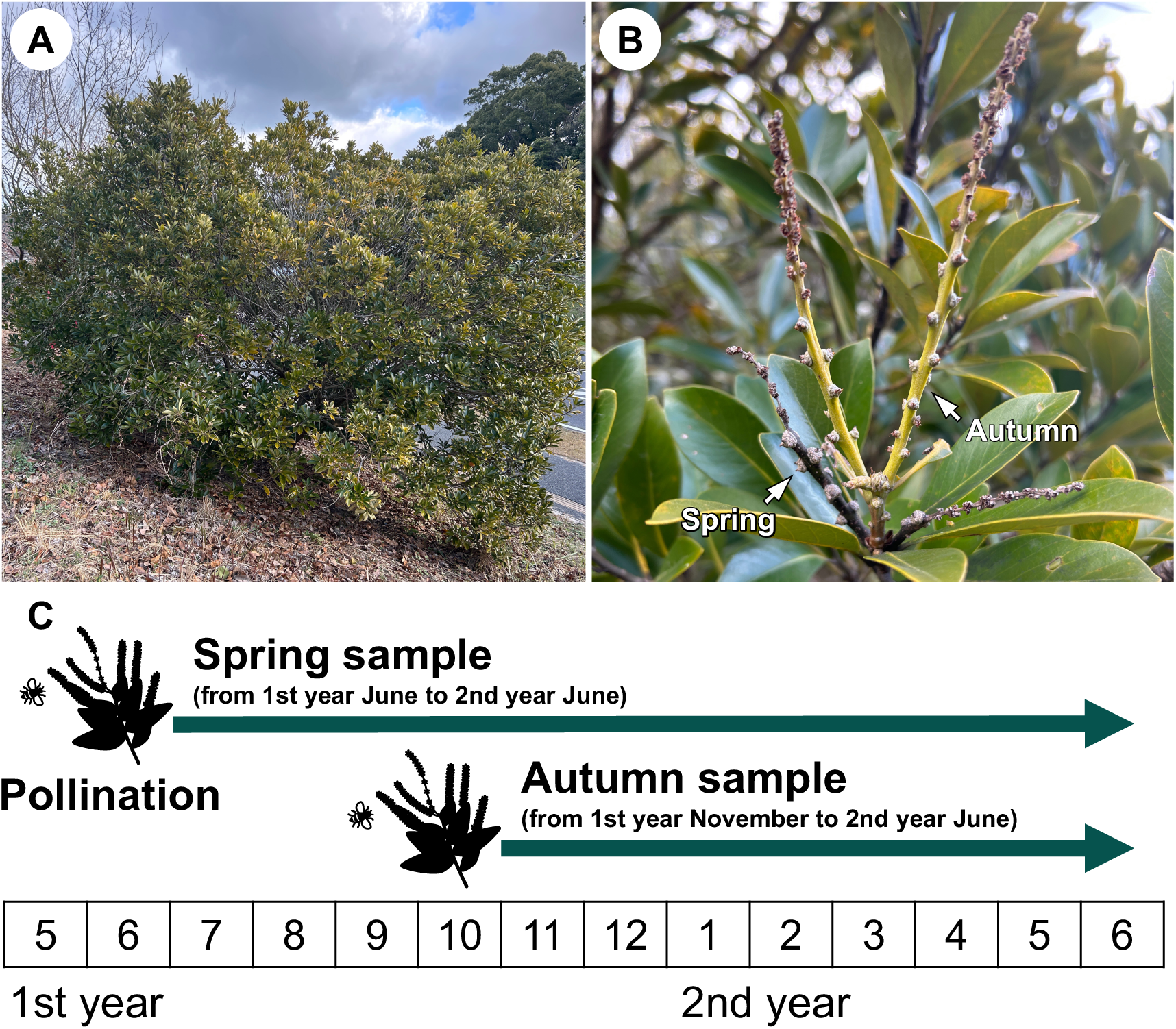
Sampling scheme of *L. edulis*. (A) One of target trees of *L. edulis* in our study. (B) A branch including both spring and autumn flowers. The photographs of (A) and (B) were taken on 24 January 2024. (C) Monthly sampling schedule. Spring samples were collected from the first year of June to the second year of June, and autumn samples were collected from the first year of October to the second year of June.

### Observations of seasonal pollen tube growth and ovule development

We employed the paraffin-embedding method to investigate the seasonal progression of the pollen tube growth and ovule development within each pistillate flower. Fixed inflorescences were dissected into each pistillate flower and then dehydrated by an ethanol series (50, 70, 80, 90, two times of 100% ethanol). After dehydration, the samples were infiltrated by the 100% xylene-ethanol series (25, 50, 75 %), and then incubated twice in 100% xylene. The samples substituted for xylene were infiltrated with 100% xylene-paraffin (Paraplast plus^®^; Sigma-Aldrich, St. Louis, MO, USA) series (25, 50, 75 %), and subsequently incubated twice in 100% paraffin. The samples were embedded in paraffin with a 57–58 °C melting point for subsequent microtome sectioning. Serial sections 6–15 μm thick were made using a rotary microtome (RX-860; Yamato Kohki Industrial Co. Ltd., Saitama, Japan). After de-waxing and rehydration, the sectioned samples were stained with 1% (w/v) toluidine blue and then stained with 0.1% aniline blue in 0.1 M K_3_PO_4_. The stained samples underwent dehydration using an ethanol series followed by a xylene series, and the slides were mounted with EUKITT^®^ (ORSA-tec, Bobingen, Germany). Pollen tubes in the pistillate flowers and the developmental stage of ovules were observed with a confocal laser-scanning microscope (FV3000; Olympus, Tokyo, Japan) using excitation lasers at 405 and 448 nm wavelengths.

Figure 2 illustrates the method for observing pollen tubes. When examining the entire section, the transmitting tissue, which does not emit a fluorescent signal, is clearly distinguished from the surrounding vascular tissue, which does emit fluorescence (Fig. 2A-C). Additionally, within the transmitting tissue, linear fluorescent signals corresponding to pollen tubes and granular fluorescent signals at the callose plugs are detected (Fig. 2D, E). Therefore, both the linear fluorescent signal representing the structure of the pollen tubes and the granular signal originating from the callose plugs observed within the transmitting tissue were used as characteristic indicators of the pollen tubes (Boavida et al. 1999; Sogo and Tobe 2006; Deng et al. 2022; Yao et al. 2023). To confirm the presence of pollen tubes detected using the method described above, we observed the pistillate flowers with the same confocal laser-scanning microscope, utilizing a combination of excitation lasers at 405 nm and 640 nm wavelengths (Appendix S1; see Supplemental Data with this article). Additionally, we classified four categories of pollen tube development based on the location of pollen tubes within the pistillate flower: style (St), style-joining site (SJS), upper ovarian locule (UOL), and micropyle (Mic). We also classified the stages of ovule development into four categories: undifferentiated (Un), ovule primordium (Op), megaspore mother cell (MMC), and mature embryo sac (MES). The number of pistillate flowers available for information on developmental stages each month in PT and ovules, including spring and autumn samples are shown in Appendix S2 (see Supplemental Data with this article). All the pistillate flowers using experiments are also shown in Appendix S3 (see Supplemental Data with this article).

**Figure 2.**
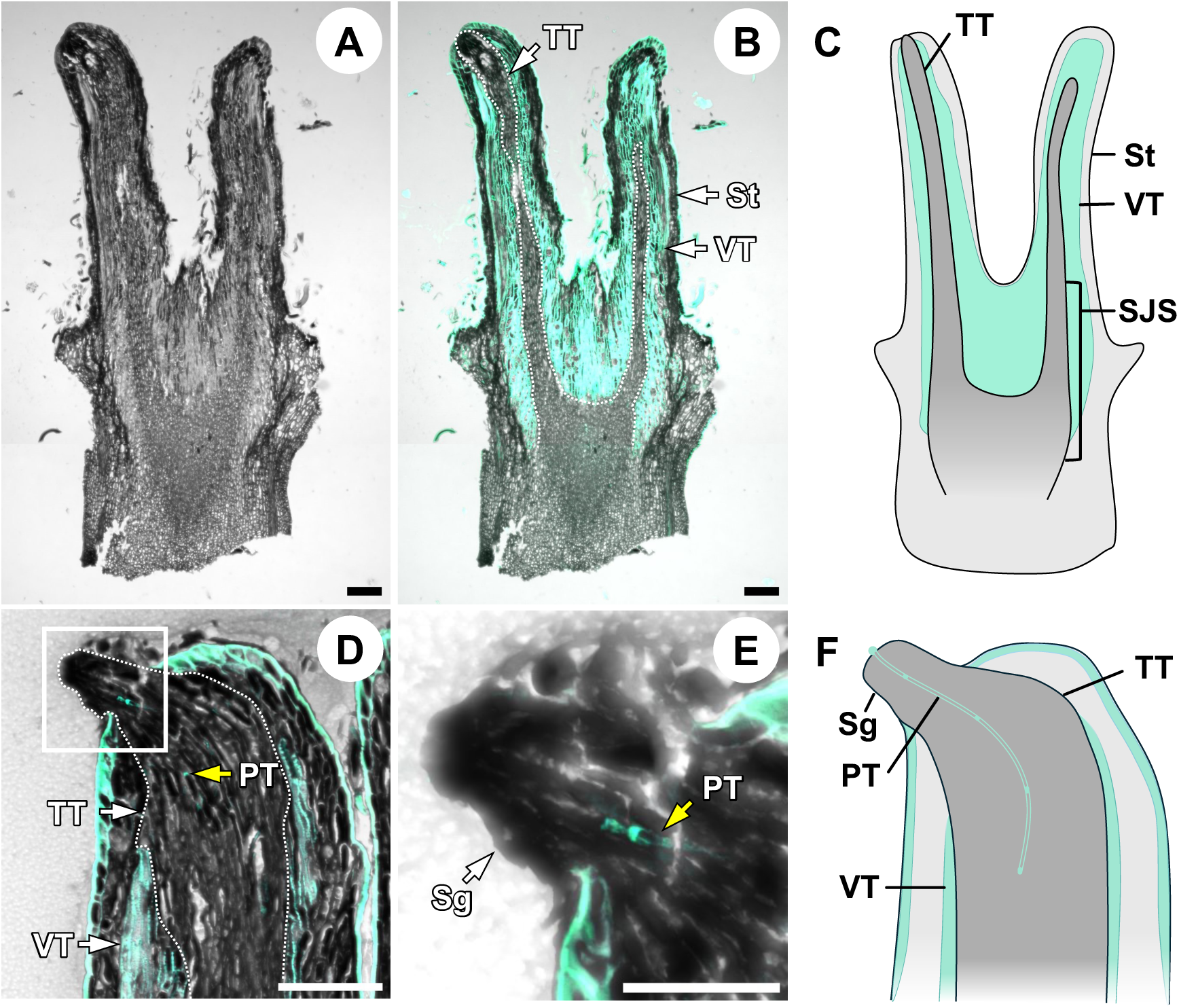
Method of pollen tube observation. (A) Overview of the entire sample. (B) Merged image of (A) with fluorescence signals. The area enclosed by the dashed line indicates the transmitting tissue. While the transmitting tissue does not emit fluorescence, a prominent fluorescent signal can be seen from the surrounding vascular tissue. (C) Schematic representation of the entire sample. The region below the style joining site (SJS), including the transmitting tissue, is shown. (D) The upper part of the style includes the stigma. The stigma is covered with a fluorescently emitting cell layer, and fluorescent signals from pollen tubes can be observed within the transmitting tissue. (E) Enlarged view of the stigma. A callose plug can be observed within the tubular pollen tube. (F) Schematic representation of the upper part of the style, including the stigma. (A) and (B) represent spring-flowering female flowers collected on 12 June 2023, and (C) and (D) represent autumn-flowering female flowers collected on 10 April 2023, respectively. The yellow arrows indicate the position of the callose plugs. PT: pollen tube; Sg: stigma; SJS: style joining site; St: style; TT: transmitting tissue; VT: vascular tissue. Scale bars = 100μm (A, B, D), 50μm (C).

## RESULTS

### Histological comparison of pollen tube growth between spring and autumn pistillate flowers

To test the overwintering-hypothesis, we compared pollen tube growth between the spring and autumn samples. In the spring samples collected right after anthesis in mid-June, most pollen tubes were observed in the transmitting tissue of the upper and middle parts of the styles (Fig. 3A). In some samples, a few pollen tubes reached the lower part of the junction of the styles (the style joining site; Fig. 3A). From July of the first year to April of the following year, the tips of the pollen tubes ceased growth and arrested at the style joining site (Fig. 3B-D). Pollen tube signals were only observed in the dark-gray stained tissues, not in the lighter gray tissues below them (Fig. 3B-D). In May of the second year, some pollen tubes resumed growth and penetrated the transmitting tissue in the upper ovarian locules (Fig. 3E). During this period, the lower part of the pistillate flower, including the ovarian wall and ovules, developed rapidly. Samples from June of the second year showed that parts of the pollen tubes were located near the ovules, with some pollen tubes positioned close to the micropyles (Fig. 3F).

**Figure 3.**
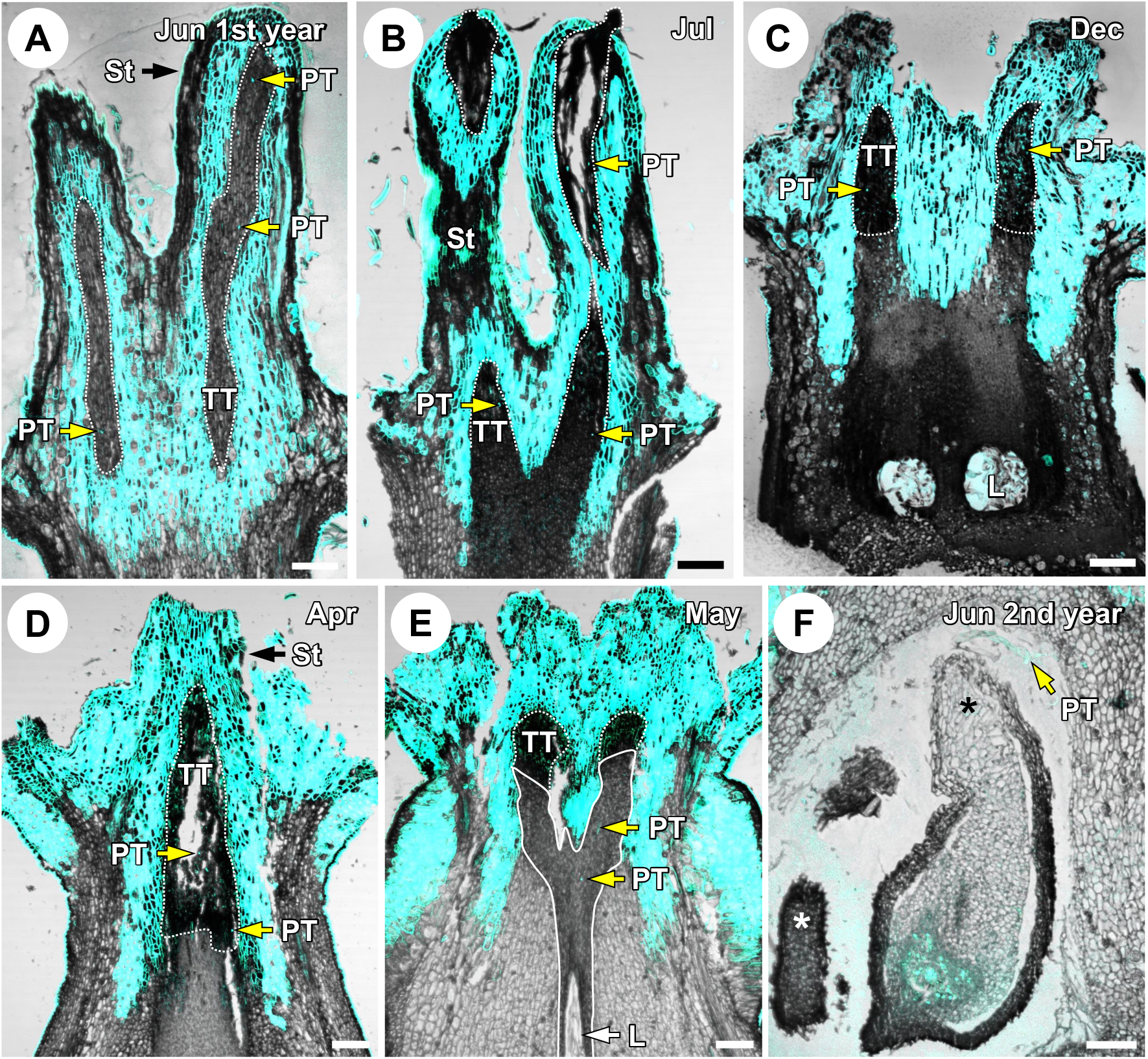
Seasonal progression of pollen tube growth in spring samples. (A) The female flower collected in June of the first year showed that some pollen tubes grew into styles (St). Some pollen tubes had reached the transmitting tissue at the base of styles. (B-D) Pollen tubes were arrested in the transmitting tissue at the style joining site (SJS) from July of the first year to April of the second year. (E) Some pollen tubes resumed elongation from the style joining site and reached the transmitting tissue of the upper ovarian locule (UOL). (F) The female flower of the second year showed that some parts of the pollen tubes were located near the micropyle (Mic). The area enclosed by a dotted line represents the transmitting tissue, and the area enclosed by a solid line indicates newly formed transmitting tissue observed between the style joining site and the ovarian locule after May of the second year. The granular or linear fluorescent signals observed within the transmitting tissue or ovule were identified as pollen tubes, while fluorescent signals observed in other areas were determined to originate from tissues other than pollen tubes. The longitudinal sections of the pistillate flowers were collected on (A) 12 Jun 2023, (B) 11 Jul 2023, (C) 9 Dec 2022, (D) 10 Apr 2023, (E) 14 May 2023, and (F) 12 Jun 2023, respectively. The locations of the ovules are marked by asterisks. PT: pollen tube; St: style; Sg: stigma; TT: transmitting tissue; L: ovarian locule. Scale bars = 100μm.

Anthesis for the autumn samples was observed from September to mid-October. In these samples, pollen tubes were arrested in the transmitting tissue of the style joining site from November of the first year to April of the second year (Fig. 4A, B). In May of the second year, some pollen tubes resumed elongation from the style joining site and were located at upper ovarian locules (Fig. 4C). In June, they reached near the ovule and micropyle (Fig. 4D). Our results demonstrate that fertilization is delayed by 13 months, from June to the following June, in the spring samples, whereas the delay was 9 months, from September to June, in the autumn samples.

**Figure 4.**
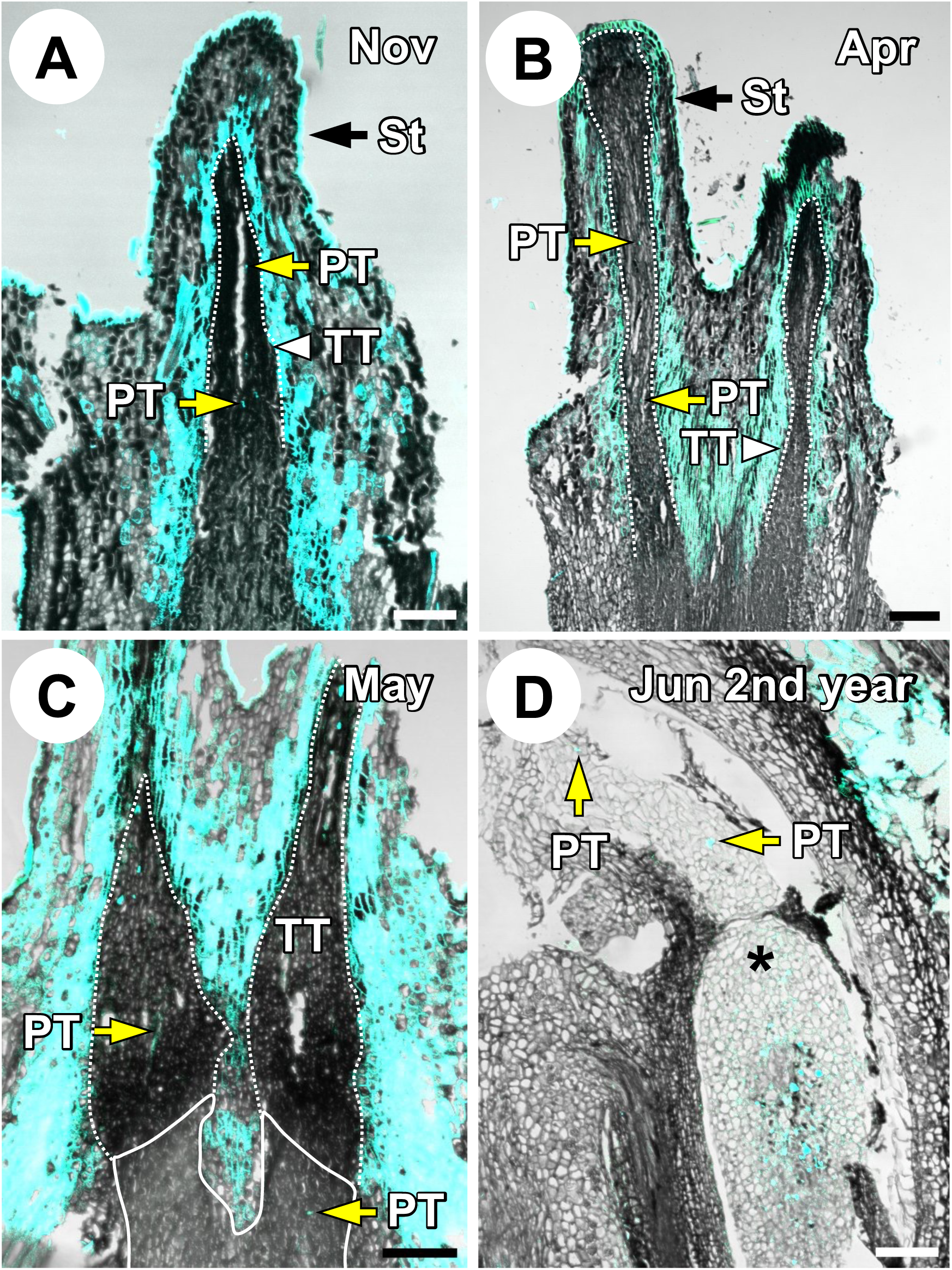
Seasonal progression of pollen tube growth in autumn samples. (A-B) Pollen tubes were arrested in the transmitting tissue at the style joining site (SJS) from November of the first year to April of the second year. (C) Some pollen tubes resumed elongation from the style joining site and reached the tissue of the upper ovarian locule (UOL). (D) The female flower of the second year showed that some parts of the pollen tubes were located near the micropyle (Mic). The area enclosed by a dotted line represents the transmitting tissue, and the area enclosed by a solid line indicates newly formed transmitting tissue observed between the style joining site and the ovarian locule after May of the second year. The granular or linear fluorescent signals observed within the transmitting tissue or ovule were identified as pollen tubes, while fluorescent signals observed in other areas were determined to originate from tissues other than pollen tubes. The longitudinal sections of the pistillate flowers were collected on (A) 11 Nov 2022, (B) 10 Apr 2023, (C) 14 May 2023, and (D) 12 Jun 2023, respectively. The location of the ovule is marked by an asterisk. PT: pollen tube; St: style; TT: transmitting tissue. Scale bars = 100μm.

### Histological comparison of ovule development between spring and autumn pistillate flowers

Seasonal ovule development in longitudinal and transverse sections of spring samples showed that in June of the first year, small and inconspicuous ovarian locules were observed at the base of the pistillate flower, but the ovule meristem was undifferentiated (Fig. 5A; Fig. 6A). From July of the first year to January of the second year, we observed that the ovarian locules enlarged, but the ovule meristem remained undeveloped (Fig. 5B, C; Fig. 6B, C). However, One out of seven samples collected in November showed ovule primordia (Fig. 5D; Appendix 2). The samples from February to April of the second year showed ovule primordia (Fig. 5E, F; Fig. 6D, E). In May of the second year, we observed that the ovarian locules further enlarged, and the ovule primordia differentiated into megaspore mother cells with outer and inner integuments (Fig. 5G; Fig. 6E). From the samples collected in June of the second year, we observed further developed ovules with mature embryo sacs formed (Fig. 5H; Fig. 6G).

**Figure 5.**
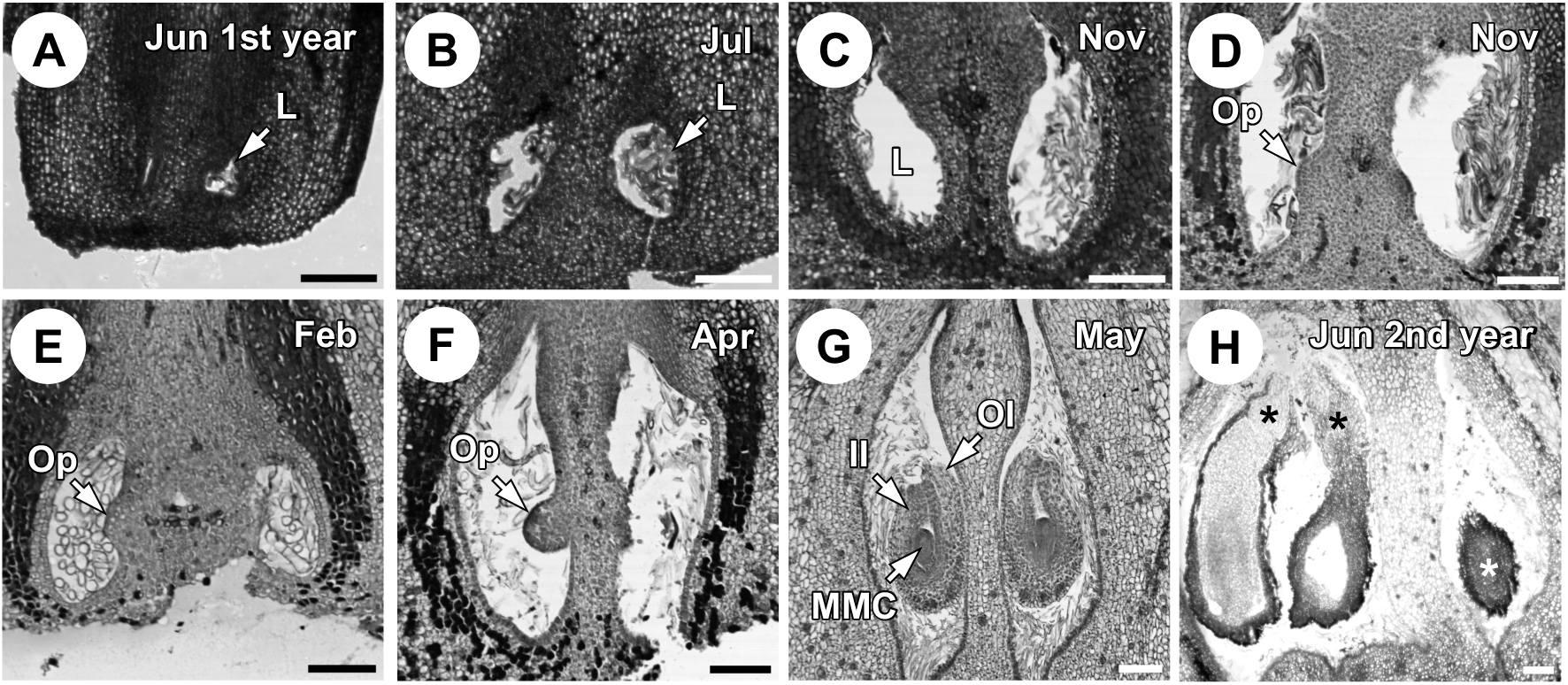
Seasonal progression of ovule development in the longitudinal section of spring samples. (A) Undeveloped ovarian locules (Un) were observed at the base of the pistillate flower collected in June of the first year. (B-C) Ovarian locules were enlarged, but the ovarian meristem had not yet developed. (D) However, some pistillate flowers collected in November showed ovule primordia (Op) in November of the first year. (E-F) Ovule primordia were observed on the placenta from February to April of the second year. (G) The ovule developed into the megaspore mother cell (MMC) stage. Outer integument and inner integument were also differentiated in this stage in May of the second year. (H) In June of the second year, the ovule developed into the matured embryo sac (MES) stage, reaching the phase when fertilization occurred. The longitudinal sections of the pistillate flowers were collected on (A) 12 Jun 2023, (B) 11 Jul 2023, (C-D) 11 Nov 2022, (E) 11 Feb 2023, (F) 10 Apr 2023, (G) 14 May 2023, and (H) 12 Jun 2023, respectively. The locations of the ovules are marked by asterisks. L: ovarian locule; Op: ovule primordium; MMC: megaspore mother cell; OI: outer integument; II: inter integument. Scale bars = 100μm.

**Figure 6.**
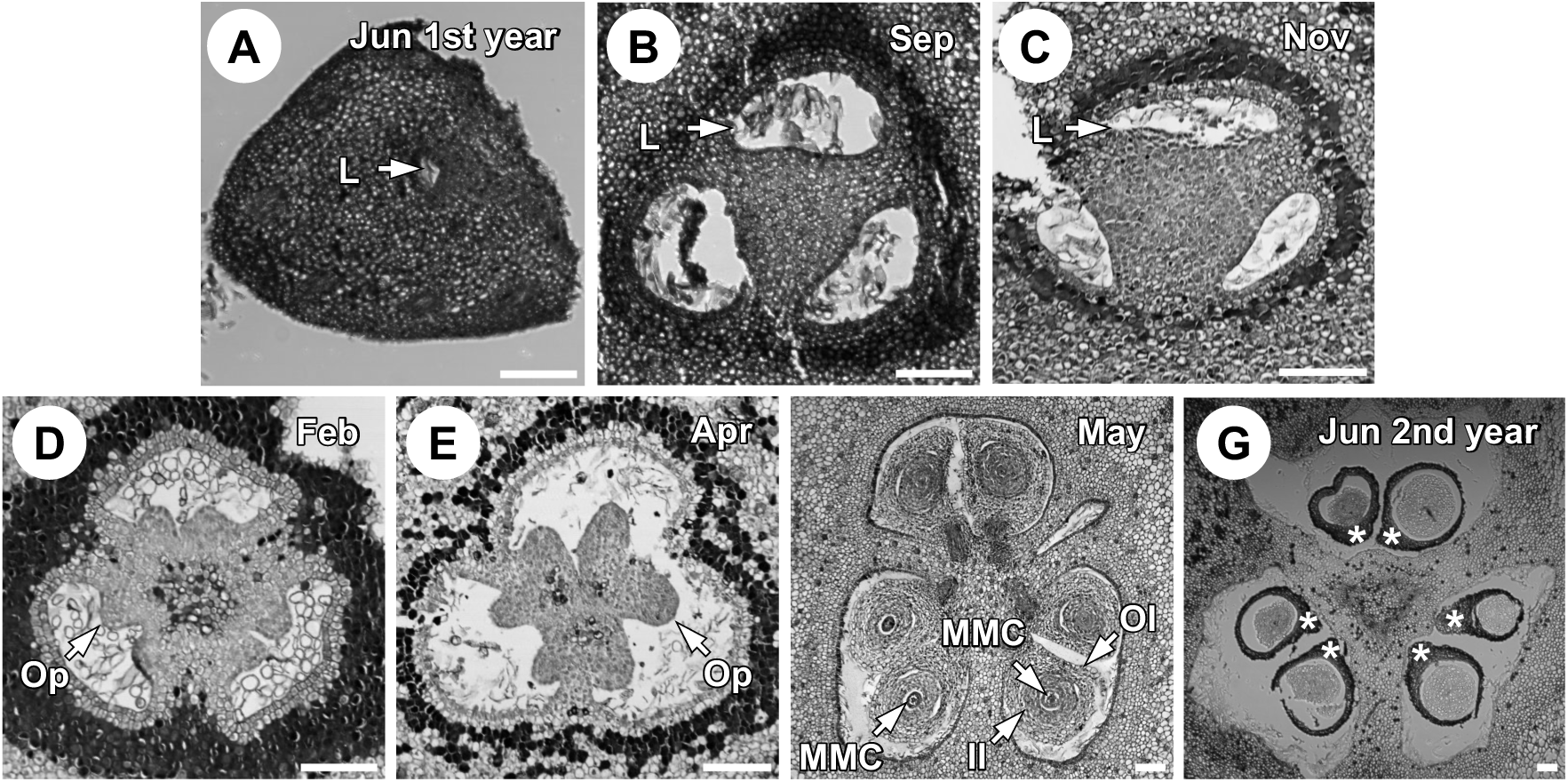
Seasonal progression of ovule development in the transverse section of spring samples. (A) Undeveloped ovarian locules (Un) were observed from the pistillate flower collected in June of the first year. (B-C) Ovarian locules were enlarged, but the ovarian meristem had not yet developed. (D-E) Ovule primordia (Op) were observed along with the placenta. (F) Ovules were differentiated at the megaspore mother cell (MMC) stage in May of the second year. (G) Ovules were developed at the mature embryo sac (MES) stage in June of the second year. The transverse sections of the pistillate flowers were collected on (A) 12 Jun 2023, (B) 20 Sep 2023, (C) 11 Nov 2022, (D) 11 Feb 2023, (E) 10 Apr 2023, (F) 14 May 2023, and (G) 12 Jun 2023, respectively. The locations of the ovules are marked by asterisks. L: ovarian locule; Op: ovule primordium; MMC: megaspore mother cell; OI: outer integument; II: inter integument. Scale bars = 100μm.

From the autumn samples collected from November of the first year to April of the second year, ovarian locules were observed, but ovule primordium was not found (Fig. 7A, B; Fig. 8A, B). In May, the lower part of the pistillate flowers including the ovary began growing rapidly, and megaspore mother cell was developed (Fig. 7C; Fig. 8C). From the samples obtained in June, we observed further developed ovules (Fig. 7D; Fig. 8D). These results indicate that ovule development begins after winter, regardless of the flowering time.

**Figure 7.**
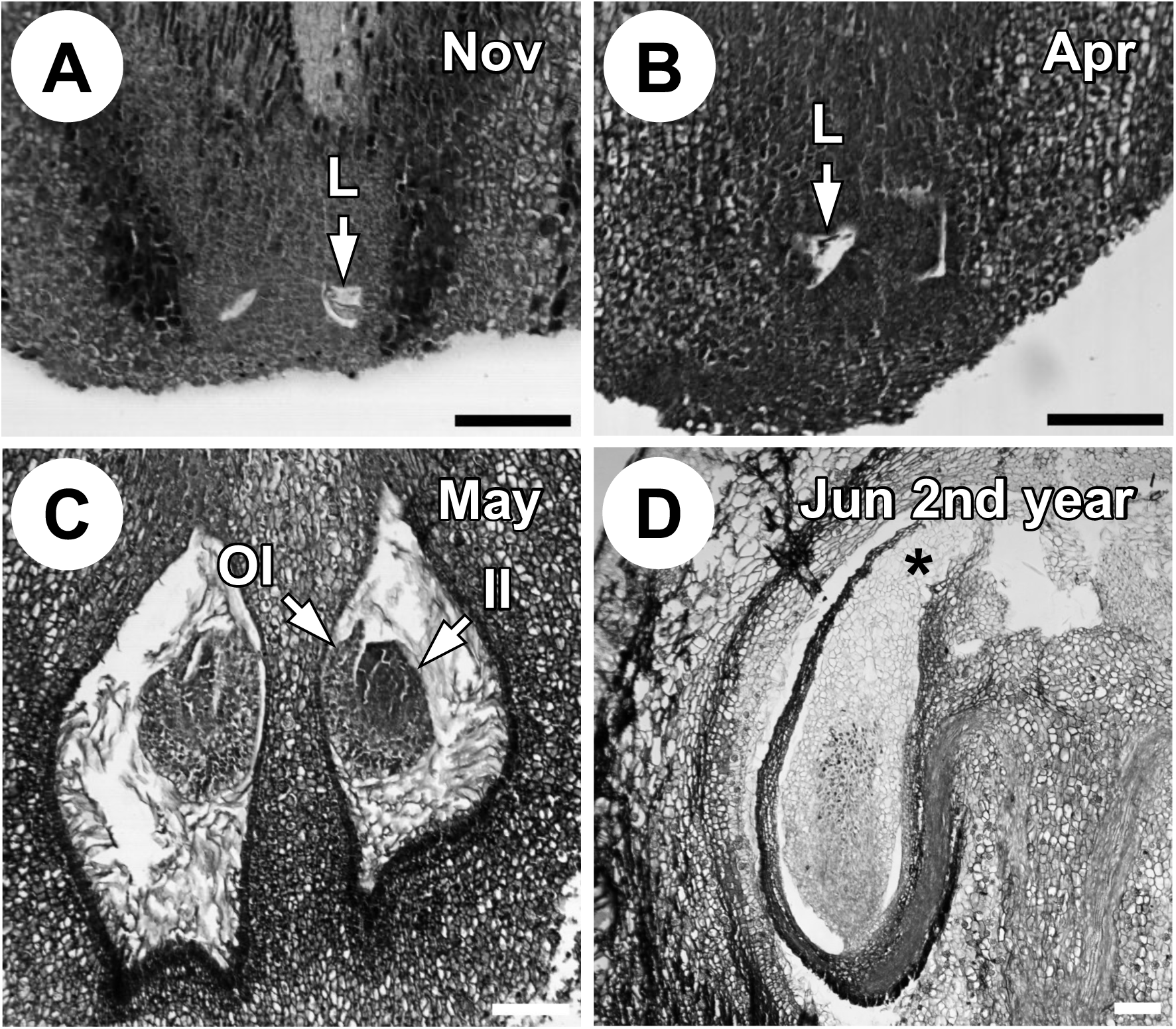
Seasonal progression of ovule development in the longitudinal section of autumn samples. (A-B) Undeveloped ovarian locules (Un) were observed at the base of the pistillate flower collected from November of the first year to April of the second year. (C) The ovule developed into the megaspore mother cell (MMC) stage in May of the second year. (D) Ovules were developed at the mature embryo sac (MES) stage in June of the second year. The longitudinal sections of the pistillate flowers were collected on (A) 11 Nov 2022, (B) 10 Apr 2023, (C) 14 May 2023, and (D) 12 Jun 2023, respectively. The locations of the ovules are marked by asterisks. L: ovarian locule; OI: outer integument; II: inter integument. Scale bars = 100μm.

**Figure 8.**
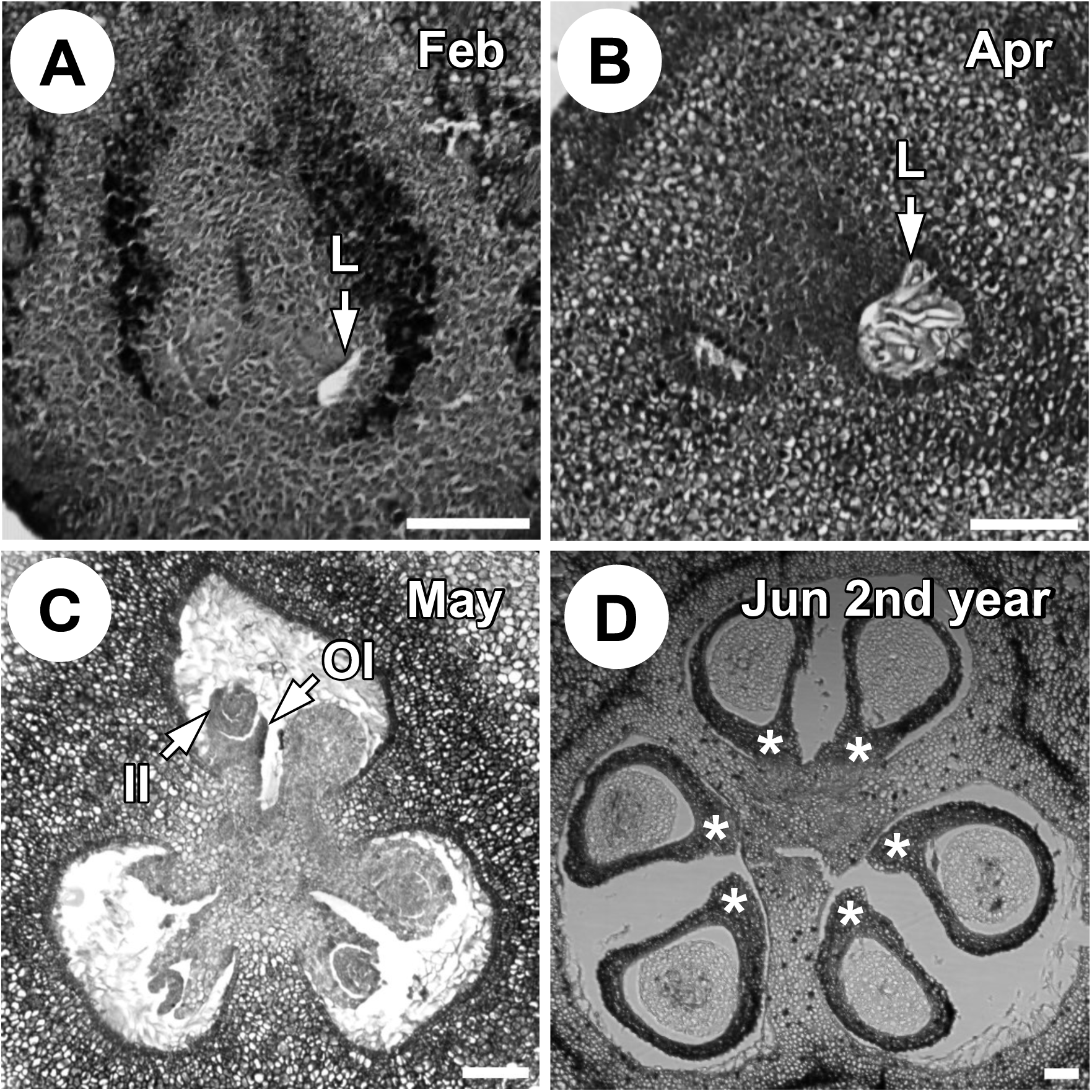
Seasonal progression of ovule development in the transverse section of autumn samples. (A-B) Undeveloped ovarian locules (Un) were observed at the base of the pistillate flower collected from November of the first year to April of the second year. © Ovules were differentiated at the megaspore mother cell (MMC) stage in May of the second year. (D) Ovules were developed at the mature embryo sac (MES) stage in June of the second year. The transverse sections of the pistillate flowers were collected on (A) 11 Feb 2023, (B) 10 Apr 2023, (C) 14 May 2023, and (D) 12 Jun 2023, respectively. The locations of the ovules are marked by asterisks. L: ovarian locule; OI: outer integument; II: inter integument. Scale bars = 100μm.

### Synchrony in the timing of pollen tube regrowth and ovule maturation

We compare the seasonal progression of pollen tube and ovule development by plotting the proportion of each category mentioned in the material and methods. For pollen tubes, the proportion in the St category remained high until April of the year following flowering, while the proportion in the SJS category increased in May of the second year (Fig. 9A, B). The transition from the St to the SJS category in May coincided with the transition of ovules from the Un or Op category to the MMC category (Fig. 7C, D). By June, the proportion of pollen tubes in the Mic category increased (Fig. 9A, B), which synchronized with the increase of ovules in the MES stage. (Fig. 9C, D).

**Figure 9.**
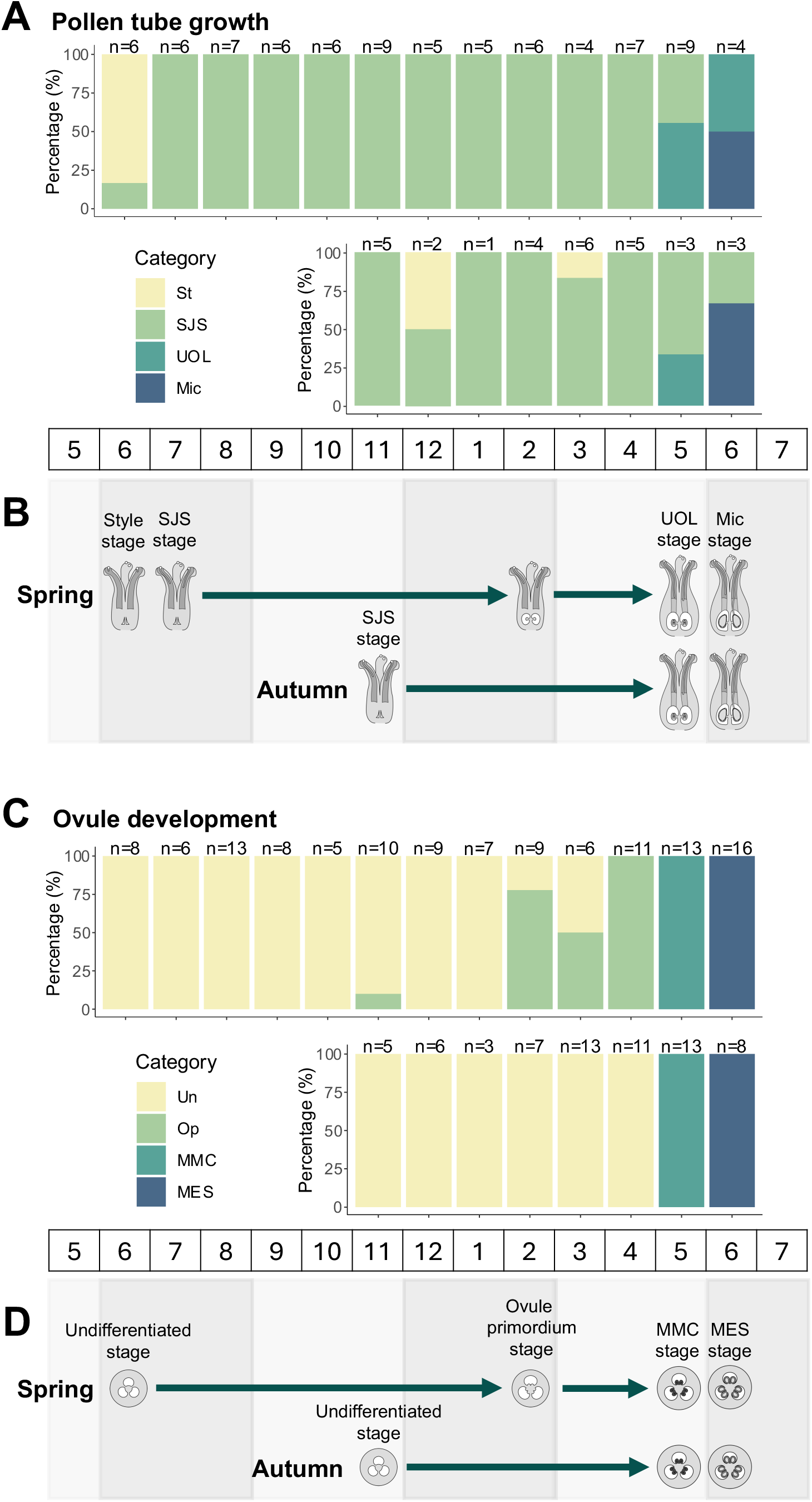
Summary of seasonal progression of pollen tube growth and ovule development. (A) Seasonal dynamics of pollen tube growth. The state of pollen tube growth is identified as four stages, style (St), style joining site (SJS), upper ovarian locule (UOL), and the micropyle (Mic) stages. The status of each month is shown as a percentage of the above four stages. The top panel shows results for spring samples, while the bottom panel shows results for autumn samples. The number of samples referenced is indicated at the top of each bar plot. (B) Summary of seasonal dynamics of pollen tube growth of *L. edulis* based on the results of (A). C Seasonal progression of ovule development. The state of ovule development is identified as four stages, undifferentiated (Un), ovule primordium (Op), megaspore mother cell (MMC), and mature embryo sac (MES) stages. The status of each month is shown as a percentage of the above four stages. The top panel shows results for spring samples, while the bottom panel shows results for autumn samples. (D) Summary of seasonal progression of ovule development of *L. edulis* based on the results of (C).

## DISCUSSION

Our results show that pistils produced in both spring and autumn overwintered and exhibit synchronous ovule fertilization after winter, which support the overwintering hypothesis. We demonstrated that pollen tubes were arrested at the style-joining site for both spring and autumn samples before winter, with their regrowth synchronized the following spring. Ovule development was delayed by 10 months in spring flowers and by 6 months in autumn flowers, resulting in a synchronized onset of ovule maturation in both sets of flowers. These findings suggest that delayed fertilization could have evolved to ensure that fertilization and seed maturation occur during the more favorable season from spring to autumn, supporting the overwintering hypothesis.

Previous studies in *Quercus* and *Lithocarpus* have demonstrated that the style joining site as a significant location for pollen tube cessation (Cecich, 1997; Boavida et al., 1999; Deng et al., 2022; Yao et al., 2023). The transmitting tissue at this site was stained deeply darkish gray, and pollen tubes are retained within this area, unable to reach the lighter gray stained tissue located below it (Deng et al., 2022). Similar to previous studies, in *L. edulis*, we observed that pollen tubes stopped growth at a position corresponding to the darkly stained transmitting tissue of the style joining site, corroborating these previous findings. Therefore, the deeply dark-stained transmitting tissue within the style joining site is a common feature associated with the arrest of pollen tube growth in Fagaceae.

After the arrest of pollen tube growth during winter, we observed that some pollen tubes resumed elongation in spring of the following year, a phenomenon also noted in *L. dealbatus* in China (Yao et al., 2023). In *L. dealbatus*, after ceasing of elongation at the upper septum and micropyle for 1-2 weeks to one month, pollen tubes resume growth, leading to fertilization (Yao et al., 2023). Although this study did not capture the exact moment of fertilization due to low pollination rates and a limited number of available samples, it is reasonable to infer that fertilization occurred in June, as indicated by embryo development observed in July (also reported in Satake et al. 2023).

Our demonstration of the over-wintering hypothesis for delayed fertilization may be applicable to other species with diverse timing of pollination and fertilization. We summarized and compared the timing of pollination and fertilization for 16 Fagaceae species from four genera reported in previous studies (*Q. gambelii* (Brown and Mogensen, 1972), *Q. alba* (Cecich, 1997; Mogensen, 1965), *Q. coccinea* (Mogensen, 1965), *Q. rubra* (Cecich, 1997), *Q. velutina* (Cecich, 1997), *Q. acutissima* (Borgardt and Nixon, 2003; Deng et al., 2022), *Q. suber* (Boavida et al., 1999; Diaz-Fernandez et al., 2004; da Costa et al., 2007), *Q. schottkyana* (Deng et al., 2008), *Q. glauca* (Satake et al., 2023), *Lithocarpus dealbatus* (Yao et al., 2023), *L. edulis* (Spring: Satake et al. 2023; this study), *L. edulis* (Autumn: Satake et al., 2023; this study), *Castanea crenata* (Nakamura, 1986; Nakamura, 1991; Nakamura, 1992; Nakamura, 1994; Nakamura, 2001; Nakamura, 2003), *C. henryi* (Fan et al., 2015), *C. mollissima* (Xiong et al., 2019; Zou et al., 2014), *C. savita* (Botta et al., 1995), and *Fagus japonica* (Sogo and Tobe, 2006); Fig. 10). The 16 species exhibit different fruiting patterns: 1-year and 2-year cycles. In 1-year fruiting species, flowering, fertilization, and fruiting all occur within the same calendar year. Conversely, in 2-year fruiting species, such as *L. edulis*, fertilization and fruiting take place in the year following flowering as shown in *L. edulis*. Regardless of the genus, pollination mode (wind or animal pollinations), and fruiting patterns (one-year fruiting or two-year fruiting), fertilization time is synchronized during spring to early summer. Among the species summarized in Fig. 10, all 1-year fruiting species flowered from April to June, followed by a fertilization delay of approximately two weeks to two months. On the other hand, in 2-year fruiting species, flowering occurred diversely from April to October, followed by a fertilization delay of 8 to 14 months, with fertilization occurring the following year, specifically in June to July of the second year. These data support the overwintering hypothesis for delayed fertilization, suggesting that the fertilization in appropriate seasons would contribute to the synchronized fruiting events in autumn (Satake and Kelly, 2021a).

**Figure 10.**
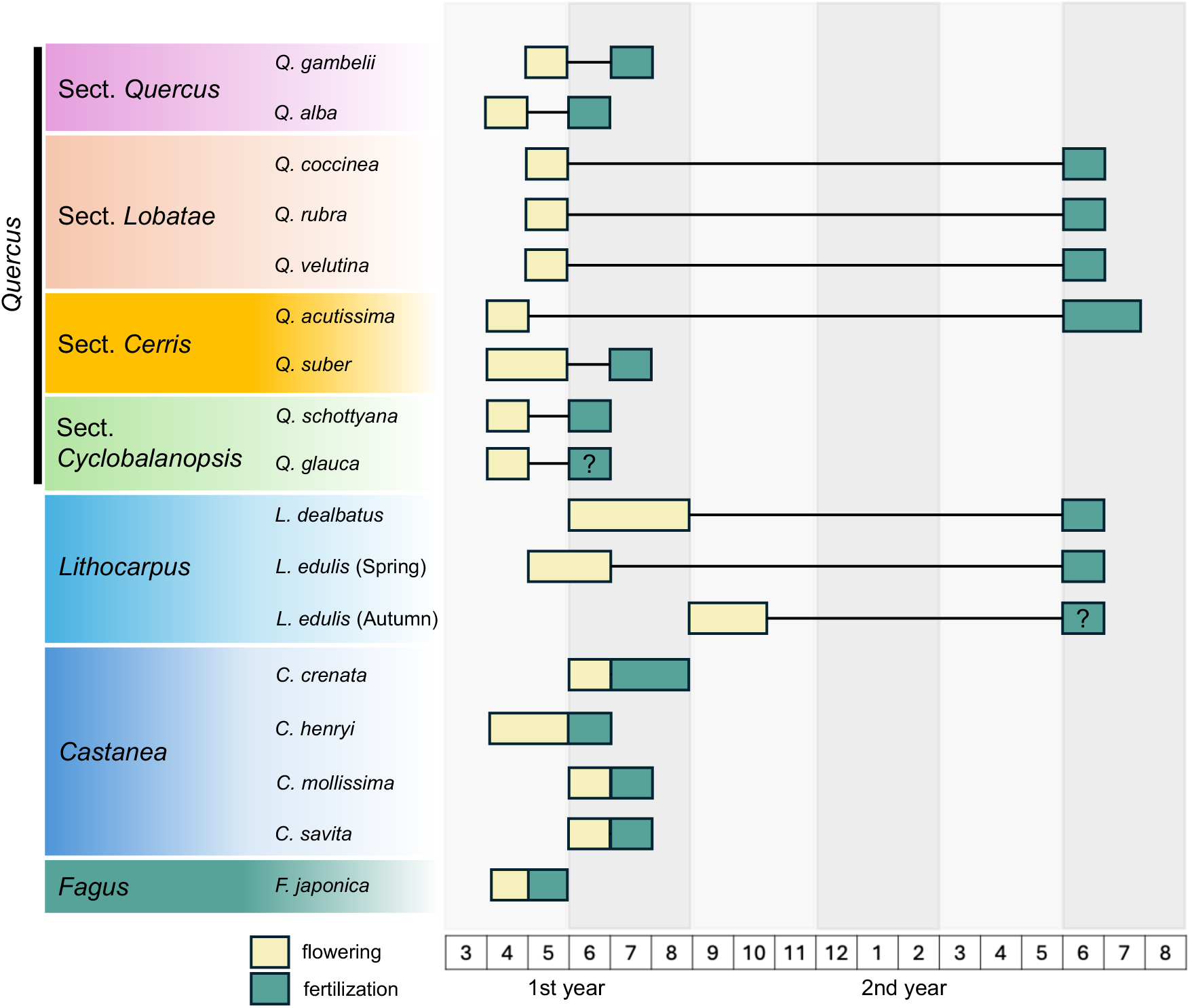
Flowering and fertilization phenology of the Fagaceae species. Yellow and green indicate the month flowering and fertilization were confirmed, respectively. The month that fertilization is presumed to have occurred is marked with “?”. This figure was compiled concerning the following research: *Q. gambelii* (Brown and Mogensen, 1972), *Q. alba* (Cecich, 1997; Mogensen, 1965), *Q. coccinea* (Mogensen, 1965), *Q. rubra* (Cecich, 1997), *Q. velutina* (Cecich, 1997), *Q. acutissima* (Borgardt and Nixon, 2003; Deng et al., 2022), *Q. suber* (Boavida et al., 1999; Diaz-Fernandez et al., 2004; da Costa et al., 2007), *Q. schottkyana* (Deng et al., 2008), *Q. glauca* (Satake et al., 2023), *Lithocarpus dealbatus* (Yao et al., 2023), *L. edulis* (Spring: Satake et al. 2023; this study), *L. edulis* (Autumn: Satake et al., 2023; this study), *Castanea crenata* (Nakamura, 1986; Nakamura, 1991; Nakamura, 1992; Nakamura, 1994; Nakamura, 2001; Nakamura, 2003), *C. henryi* (Fan et al., 2015), *C. mollissima* (Xiong et al., 2019; Zou et al., 2014), *C. savita* (Botta et al., 1995), and *Fagus japonica* (Sogo and Tobe, 2006). In *Q. suber*, 1-year fruiting tree, 2-year fruiting tree, and a mixture of both types in the same individual have been reported (Elena-Rossello et al., 1993; Diaz-Fernandez et al., 2004). In this study, only the results of the 1-year fruiting type, for which results have already been published, are shown.

To explore the molecular mechanism of delayed fertilization in response to winter cold, employing genome-wide transcriptomics analyses under seasonal conditions (Satake et al. 2022), alongside integrated analyses of gene expression patterns related to ovule development, pollen tube growth, and fertilization, combined with the histological studies reported here, will be a promising approach to elucidate the molecular mechanisms and evolutionary aspects of delayed fertilization.

## CONCLUSIONS

We examined the location of pollen tubes and the developmental stage of ovules in female flowers produced in spring or autumn nearly every month from pollination to fertilization in *Lithocarpus edulis*, a species exhibiting a 2-year fruiting pattern. Our findings indicate that the process leading to fertilization remains synchronous across different flowering periods, supporting the overwintering hypothesis. We observed that ovules stay undifferentiated until exposed to winter conditions, suggesting that cold temperatures may synchronize the development stages of ovules produced at different times. These results imply that delayed fertilization may have evolved as a strategy to ensure that fertilization and seed maturation coincide with favorable seasons, thereby avoiding the adversities of winter.

## Acknowledgments

The authors thank Yukiko Ogino and Kayoko Ohta for helping technical supports of the experiments. We thank Tetsukazu Yahara for helpful comments and discussion on this study. This work was supported by Grants from JSPS KAKENHI (grant no. JP21H04781) to A.S.

## Author Contributions

T.S. and A.S. conceived the research idea. T.S. collected the samples and carried out the experiments with the support of K.O., M.K., and A.S. T.S. and A.S. wrote the original draft, and all authors approved the final submission.

## Data Availability Statement

Data from this study are available in Appendices S1, S2, and S3.

## Online Supporting Information

Additional supporting information may be found online in the Supporting Information section at the end of the article, including Appendices S1 (“Seasonal developmental stages of pollen tubes and ovules in *L. edulis* for spring and autumn samples. The numbers in parentheses indicate the number of pistillate flowers for which developmental stages were identified”), S2 (“The number of pistillate flowers examined for observation in *L. edulis*. The results for each month were based on pistillate flowers obtained from one individual”), and S3 (“The detection of pollen tubes using excitation lasers with wavelengths of 405 nm and 640 nm”).

## Abbreviations

II: inner integument
L: locule
MES: mature embryo sac
Mic: micropyle
MMC: megaspore mother cell
OI: outer integument
Op: ovule primordia
PT: pollen tube
SJS: style joining site
St: style
TT: transmitting tissue
UOL: upper ovarian locule

## Supplemental materials

**Appendix S1.**
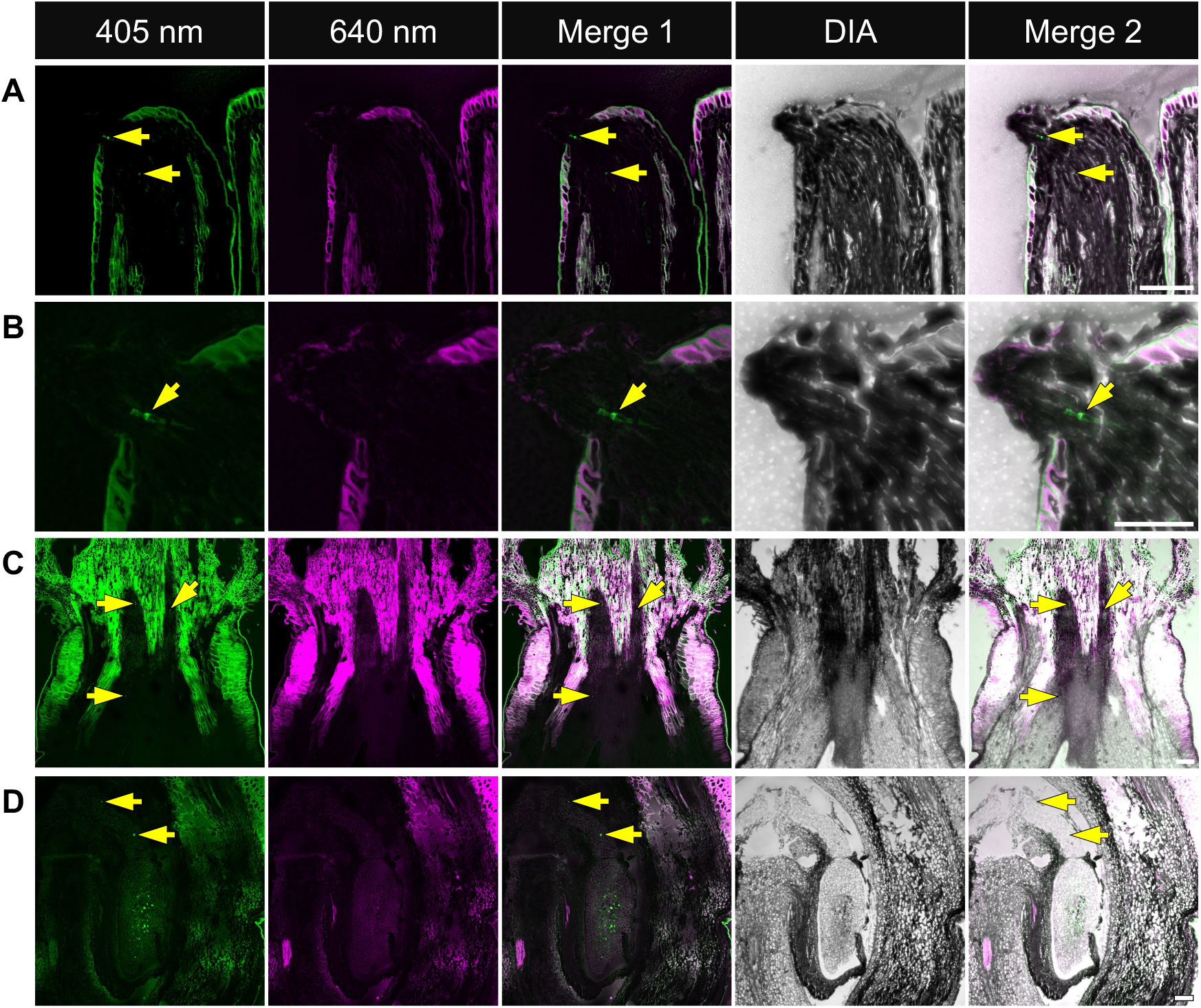
The detection of pollen tubes using excitation lasers with wavelengths of 405 nm and 640 nm. The images display five different channels: excitation at 405 nm (green), excitation at 640 nm (magenta), merged channels (405 nm + 640 nm), bright field (DIA), and final merge (405 nm + 640 nm + DIA). (A) Pollen tube signals were detected inside the style (St). (B) Magnified view of the area near the stigma in (A). (C) Pollen tube signals were detected at the style joining site (SJS) and upper ovarian locule (UOL). (D) Pollen tube signals were detected in the micropyle (Mic). The granular or linear fluorescent signals observed within the transmitting tissue or ovule were identified as pollen tubes indicated by the yellow arrows, while signals in other regions were determined to originate from non-pollen tube tissues. The longitudinal sections of the pistillate flowers were collected on (A, B) 10 Apr 2023, (C) 14 May 2023, and (D) 12 Jun 2023, respectively. Scale bars = 100μm (A, C, D), 50μm (B).

**Appendix S2.**
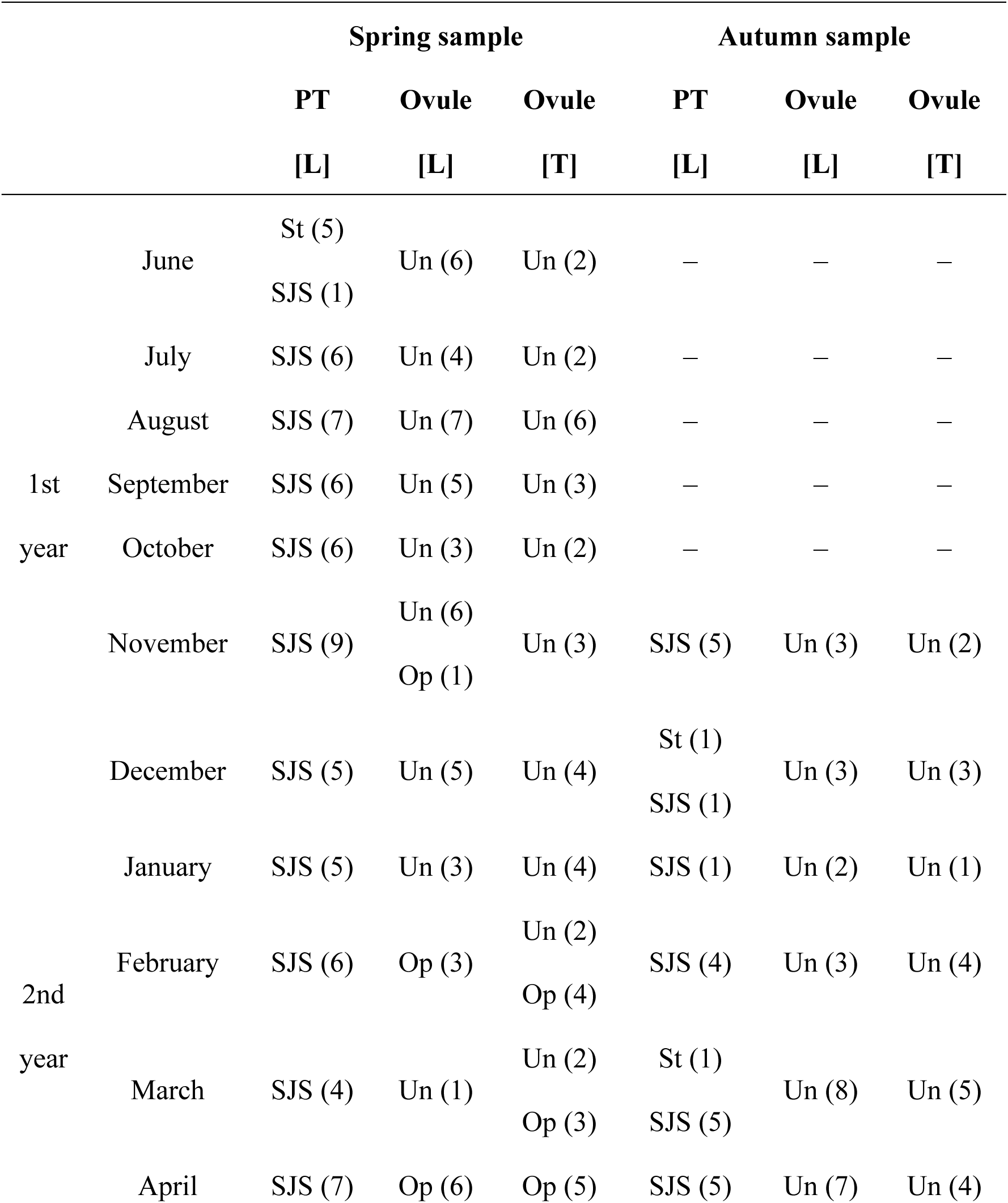
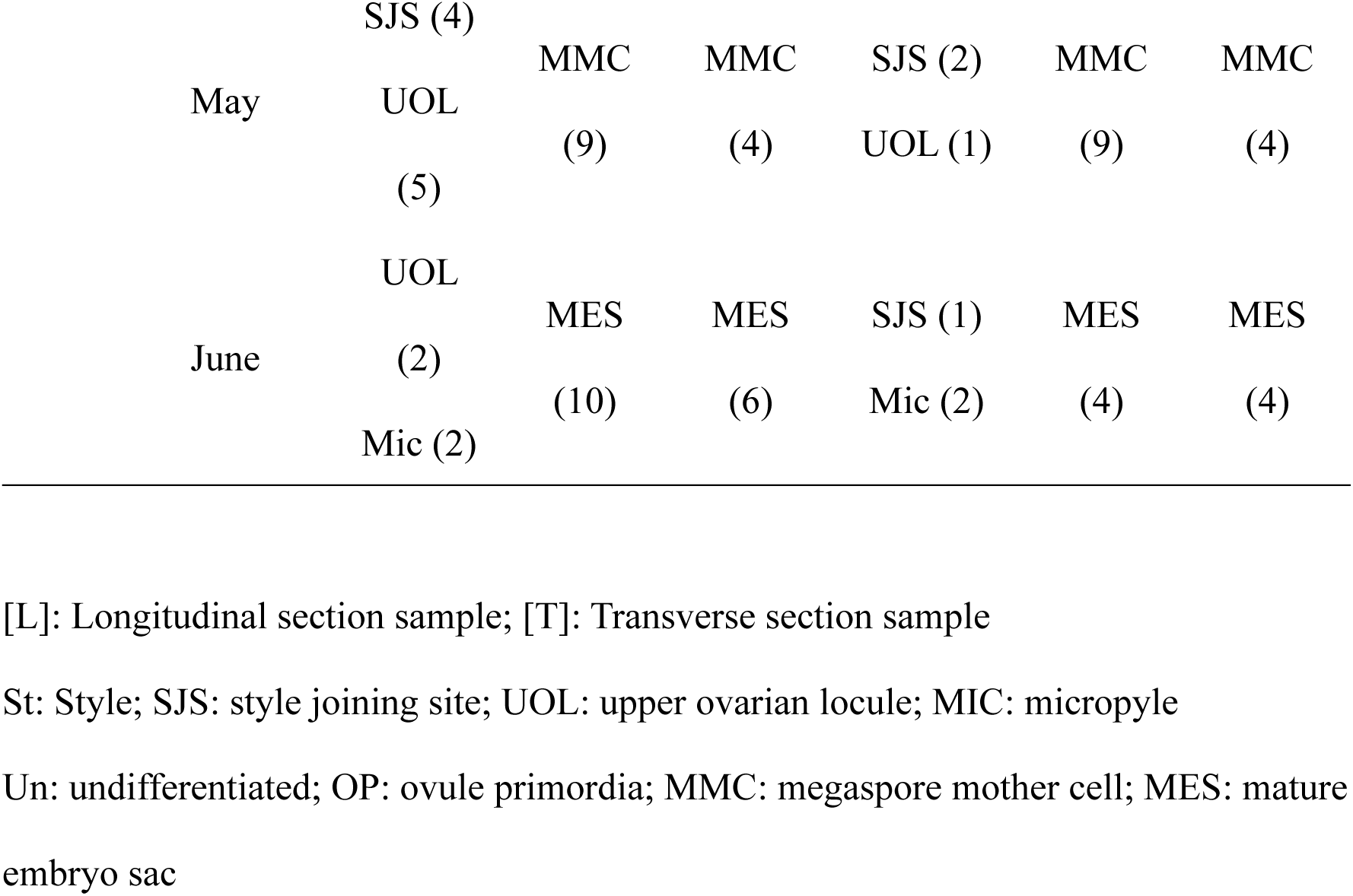
Seasonal developmental stages of pollen tubes and ovules in *L. edulis* for spring and autumn samples. The numbers in parentheses indicate the number of pistillate flowers for which developmental stages were identified.

**Appendix S3.** The number of pistillate flowers examined for observation in *L. edulis*. The results for each month were based on pistillate flowers obtained from one individual.

|  |  | Spring sample |  |  | Autumn sample |  |  |
| --- | --- | --- | --- | --- | --- | --- | --- |
|  |  | PT | Ovule | Ovule | PT | Ovule | Ovule |
|  |  | [L] | [L] | [T] | [L] | [L] | [T] |
| 1st<br>year | June | 10 | 10 | 5 | — | — | — |
|  | July | 6 | 6 | 3 | — | — | — |
|  | August | 7 | 7 | 7 | — | — | — |
|  | September | 6 | 6 | 6 | — | — | — |
|  | October | 6 | 6 | 4 | — | — | — |
|  | November | 17 | 17 | 4 | 16 | 16 | 6 |
|  | December | 6 | 6 | 4 | 8 | 8 | 7 |
| 2nd<br>year | January | 8 | 8 | 6 | 13 | 13 | 11 |
|  | February | 6 | 6 | 6 | 6 | 6 | 6 |
|  | March | 4 | 4 | 5 | 8 | 8 | 8 |
|  | April | 7 | 7 | 5 | 8 | 8 | 6 |
|  | May | 9 | 9 | 4 | 9 | 9 | 4 |
|  | June | 12 | 12 | 7 | 12 | 12 | 4 |

[L]: Longitudinal section sample; [T]: Transverse section sample

## Notes

### Competing Interest Statement

The authors have declared no competing interest.

